# Phase response curve and RNA-sequencing demonstrate spiders’ sensitivity to light and pinpoint candidate light-responsive genes

**DOI:** 10.64898/2026.02.17.706298

**Authors:** Natalia Toporikova, Wenduo Cheng, Leyuan Qian, Andrew Mah, Thomas Clarke, Thomas C. Jones, Darrell Moore, Nadia A. Ayoub

**Affiliations:** Department of Biology, Washington and Lee University, Lexington, VA, 24450, USA; Computational Biology Department, Carnegie Mellon University, Pittsburgh, PA 15213, USA; Department of Biostatistics, School of Public Health, University of Michigan, Ann Arbor, USA; Center for Neural Science, New York University, New York, New York, 10003, USA; Department of Biological Sciences, East Tennessee State University, Box 70703, Johnson City, TN, 37604, USA

**Keywords:** Circadian rhythms, *Metazygia wittfeldae*, Araneidae, entrainment, RNA sequencing, *de novo* transcriptomics, mathematical modeling, phase response curve

## Abstract

Spiders can maintain a wide range of free-running periods while still being entrained to a 24-hour day. To investigate the underlying mechanism of this entrainment, we constructed the phase-response curve (PRC) for the orb weaver, *Metazygia wittfeldae*, by subjecting the spiders to one-hour light pulses at various times throughout the circadian day. The resulting type 0 PRC showed high amplitude (> 6 hour) phase advance and delays when the light pulse was applied during circadian time (CT) 16-18, with a break point of advances to delays at CT 17. We then investigated the genetic mechanism of the phase response to light by splitting *M. wittfeldae* adult females entrained to 12 hours light:12 hours dark (LD 12:12) into two groups. One group received a 1-hour light pulse 5 hours after lights off (CT17), and one group did not. We then sacrificed spiders for RNA isolations 1 and 10 hours after the light pulse. We identified numerous genes that were downregulated by the light pulse 1 hour after the pulse relative to no pulse group. Intriguingly, many of these genes had a flipped pattern of expression 9 hours later – the pulse group had higher expression than the no pulse group. This pattern is consistent with the shifted phase of locomotor activity expected after the light pulse application. We also identified clock gene homologs in *M. wittfeldae* that had distinct expression patterns from other arthropods.

## Introduction

Daily rhythms are ubiquitous across life, from the sleep-wake cycle in humans (1) to cell division in fungi (2) to nitrogen fixation in cyanobacteria (3). These rhythms are termed circadian because they follow a natural day-night environment and continue to cycle for approximately (circa) 24 hours even in constant darkness. These internal circadian “clocks” are thought to provide a fitness advantage by allowing organisms to anticipate and prepare for daily changes in their environment (4). The proteins that control the endogenous circadian clock differ between plants, fungi, animals, and cyanobacteria, suggesting that clocks may have evolved independently multiple times, further supporting their evolutionary benefit (5).

Despite differences in individual components, all molecular clocks share a common feature: a transcription-translation negative feedback loop (6). For instance, in the fruit fly, *Drosophila melanogaster*, the clock genes *Period* (*per*) and *Timeless* (*tim*) form a negative feedback loop by encoding proteins that inhibit their own transcription. Transcription of *per* and *tim* is activated by a heterodimer of basic helix-loop-helix transcription factors, CLOCK (CLK) and CYCLE (CYC), which bind the E-box in the *per* and *tim* promoters (7). After translation, PER and TIM proteins dimerize, enter the nucleus, and inhibit CLK and CYC from binding the E-boxes. Once PER and TIM degrade, CLK and CYC can again promote transcription (8–12). This feedback loop generates daily cycles of *per* mRNA and PER protein, which in turn influence the daily cycles of locomotion and other physiological processes (13). The transcription-translation feedback loop described above is conserved across animals, with some differences: in mammals, butterflies, and many other arthropods, BMAL1 (a homolog of CYC) dimerizes with CLK to promote transcription of *per* and *Cryptochrome* genes, termed mammalian-type *cry* or *cry2* in arthropods. PER and CRY2 then dimerize and enter the nucleus to block transcription (14–16).

The environment also affects the transcription-translation feedback loop. Since the endogenous circadian period, called the free-running period (FRP), is not exactly 24 hours, external cues must entrain the clock to the earth’s rotational period (17). Light is the best described and probably the most powerful such cue, or zeitgeber (time giver (18)). In mammals, light reaching the eyes changes the structure of melanopsin, which in turn starts a cascade that eventually results in higher transcription of *per* genes (19–24). In fruit flies, light directly changes the structure of CRY1 so that it binds TIM, leading to the ubiquitination and, thus, degradation of TIM (25). Without TIM, PER becomes unstable so that it cannot block transcription of CLK and CYC, essentially resetting the clock every morning. The compound eyes also mediate light entrainment in fruit flies, but through a mechanism that is not well understood (26).

The phase response curve (PRC) is a classic method for investigating the nature of the endogenous circadian cycle and its entrainment by light (27,28). Essentially, the effect of light on the cycle differs depending on the point in the cycle (phase) at which light is applied. At some points of the cycle, light can shift the phase, while at others, it cannot (29,30). For instance, in mammals, light in the early morning advances the phase by promoting the transcription of *per* genes slightly earlier than it would naturally. On the other hand, light in the evening delays the phase by increasing transcription of *per* genes (31–33). Light pulses in the middle of the subjective day, even in a constantly dim environment, typically do not shift a cycle in humans (30,34). Similarly, fruit flies experience phase delays if pulsed with light in the early evening, when *per* and *tim* are rising. In contrast, flies experience phase advances if pulsed with light in the late evening when *per* and *tim* would be naturally decreasing (35,36).

The largest values (or amplitude) of phase advances or delays in a PRC reflect the strength of the circadian oscillator and/or its sensitivity to light (37). In wild-type flies with intact *cry1*, the PRC has much higher amplitude advances and delays than in *cry1*-mutants (38). Mutants without functional CRY1 still experience low amplitude phase shifts, presumably due to low-sensitivity compound eye-mediated light detection (26,38). A high amplitude PRC can also be indicative of a weak oscillator (39). For instance, clock mutants with low differences between the peak and trough of *per* expression have high amplitude PRCs (40).

Spiders are an emerging system for investigating the evolution of circadian rhythms and their entrainment. Thus far, circadian rhythmicity of locomotor activity has been well documented in multiple species from 6 divergent families of spiders (41–48). Anti-predator behavior in one orb-weaving species is also circadian (49), which appears to be driven by circadian cycling in octopamine levels (50). Most spiders also build their webs on a daily cycle (51), but the persistence of the rhythm in constant conditions has not yet been shown.

Surprisingly, the FRP of locomotor activity can be highly variable among and within spider species. For instance, the trashline orb weaver *Cyclosa turbinata* (Araneidae) has an average FRP of 18.7 hours in constant darkness, the shortest known period for any free-living organism. In addition, the ranges of FRPs exhibited by three cobweb weaving spider species (Theridiidae) were large: 19-23.5 h in the common house spider *Parasteatoda tepidariorum* (mean FRP = 21.7 h), 20-29 h in the subsocial spider *Anelosimus studiosus* (mean FRP = 23.1), and 20-30.1 h (mean FRP = 24.5) in the southern black widow *Latrodectus mactans* (48). Even the orb weaver *Metazygia wittfeldae* (Araneidae), which possesses an almost “typical” FRP (mean = 22.7 h) varied from 19 – 24 hours (47). The three theridiid and two araneid species have FRPs that vary an order of magnitude more than those of most animals examined thus far (48). Nevertheless, all the individuals of these species rapidly entrained to cycles of 12 hours light, 12 hours dark (LD 12:12) (46–48).

The wide range of FRPs and the robust entrainment to light by spiders suggests they may have a weak oscillator or a heightened sensitivity to light relative to other arthropods. However, very little is known about molecular clock mechanisms in spiders. We thus used a multi-pronged approach to determine the nature of the circadian cycle in one orb-weaving species, *M. wittfeldae*. First, we constructed a PRC to determine when the system is most sensitive to light and to discover the level of light response. Second, we used the results of the PRC to determine the optimal time to expose spiders to a light pulse for RNA sequencing. To further investigate how light might shift the phase of gene expression, we explored two mathematical models of clock genes. We identified a large cluster of genes in our RNA sequencing experiment that fit the expectations for phase shifting, and thus represent candidate genes controlled by the circadian clock. We also identified homologs of 6 canonical clock genes and found that two fit the expected pattern for phase shifting, while the others had markedly different patterns of expression.

## Results

### *M. wittfeldae* has a high amplitude Phase Response Curve

To our knowledge, we have derived the first PRC to light for any spider species (Figure 1a). Of 137 individual spiders, 102 survived the entire duration of the experiment and yielded phase advances or delays in response to a 1-hour light pulse. There was no significant difference (t-test for correlated samples, *P* = 0.29) between the mean FRPs before (23.49 ± 0.06 h) and after (23.43 ± 0.09 h) the light pulse, indicating the absence of light stimulus after-effects on the circadian period. Indicative of a type 0 PRC (52), there are high amplitude (> 6 hr) phase advances and phase delays and an abrupt discontinuity or “breakpoint” at the transition from delays to advances at approximately CT 17-18. In contrast, the predominant PRC found in nature is type 1, in which the phase shifts are relatively small (< 6 hr; often < 2 hr), and the transition from delays to advances along the PRC is continuous (Johnson 1992).

**Figure 1.**
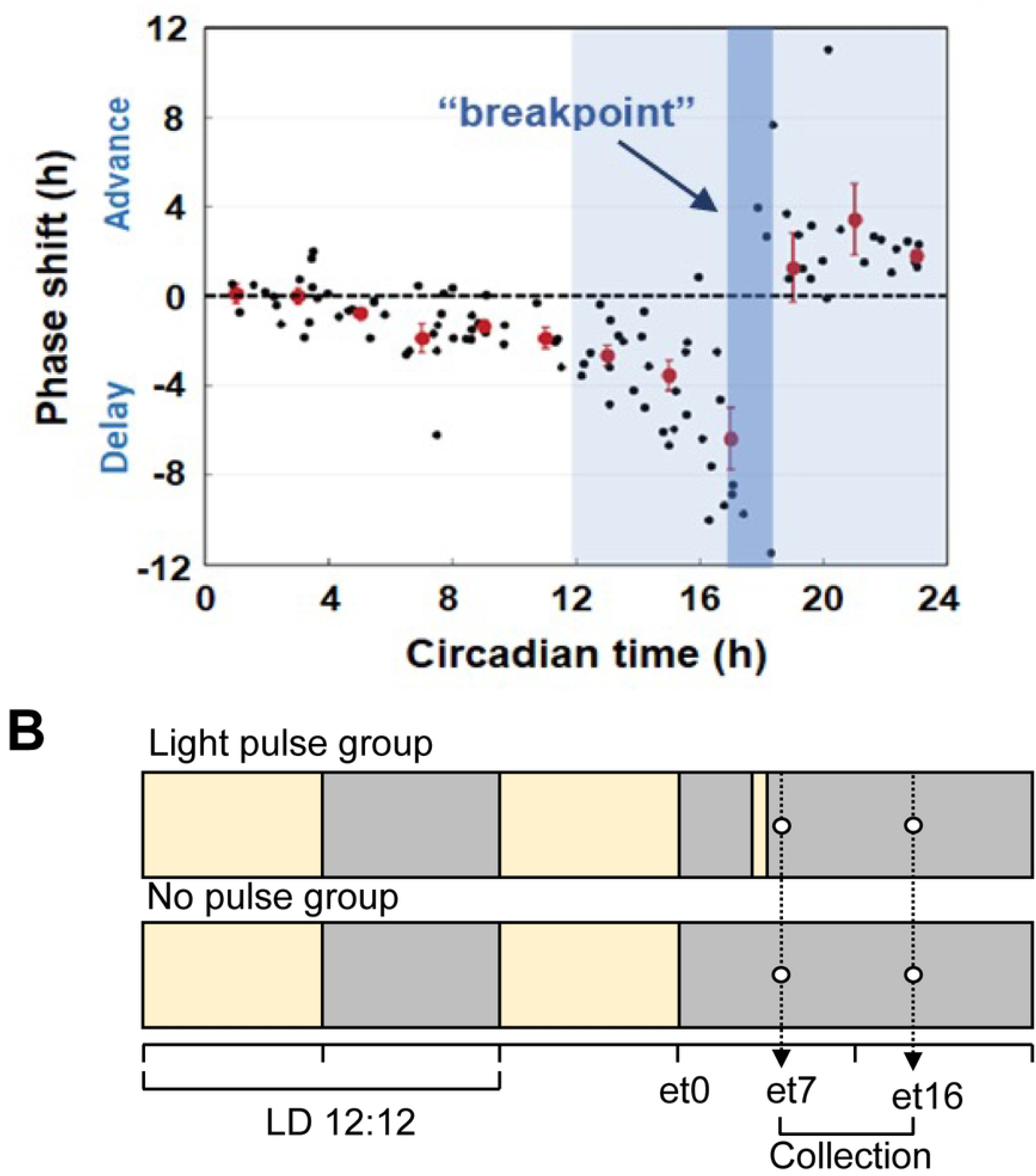
Experimental phase response curve informed gene expression experiment. (A) The phase response curve of locomotor activity of *M. wittfeldae* inferred from the application of one-hour light pulses at different phases in an individual spider’s circadian day. Each point represents the phase change induced by light at that individual’s circadian time. Red dots show mean phase shift ± SEM for 2 hour blocks (n varies by time point, total n=102). (B) Timeline for RNA-sequencing experiment based on the PRC, which indicated that a 1-hour pulse of light applied at CT 17-18 (5-6 hours after lights off) would be most likely to induce a phase shift in gene expression. Spiders were entrained to LD 12:12 for 5 days. At the end of the last light period (et0), half of the subjects were left in constant darkness (bottom plot), and the other half was exposed to 1 hour long light pulse 5 hours after darkness onset (et5). The samples were collected in both groups one hour after the end of the light pulse (et7) and 10 hours after the pulse (et16).

### PRC suggests time for maximal gene expression response to light

The PRC showed that applying a 1-hour light pulse at CT 17-18 caused maximal phase advances and delays in locomotor activity (Figure 1A). We reasoned that a 1-hour light pulse at this time should additionally shift the phase of expression for circadian genes. We thus designed an experiment for gene expression analysis in which spiders were entrained to a 12:12 LD cycle for 5 days to synchronize gene expression. On the evening of the 5^th^ day, the lights were turned off (*et0*), but one group of 10 spiders received a 1-hour pulse of light (*pulse* group) 5 hours after lights off (*et5-et6*) and the other group of 10 did not (*no pulse* group). The timing of the pulse of light is analogous to CT 17-18 (Figure 1A).

Due to resource-limitations, we could only sample for RNA-sequencing at two times after the light pulse. Theoretically, the light pulse should shift the phase of circadian genes, but the exact shift is hard to predict for a species without prior circadian genetic information. We explored how a 1-hour light pulse might shift the phase of gene expression by modifying two mathematical models (Supplementary File 1: Mathematical modeling and Figures S1-S2). Both suggested that collecting 1 hour and 10 hours after the light pulse would maximize the differential expression of circadian genes between *pulse* and *no-pulse* groups. We thus sampled 5 spiders for RNA isolations from *pulse* and *no-pulse* groups 1 hour after the end of the light pulse (*et7*) and 10 hours after the light pulse (*et16*) (Figure 1B).

Since no genomic resources exist for *M. wittfeldae*, we *de novo* assembled 266,300 transcripts from the 20 individual RNA-seq libraries, representing 184,077 Trinity-defined “genes”. Annotation of the “genes” resulted in 9 categories of sequences, 4 of which were unlikely to encode proteins and were thus removed prior to further analyses (Supplementary File 1: Supplementary Table S1). Of the retained translated genes, 25,489 were further annotated by significant alignment to a protein in SwissProt and/or PFAM, and 25,127 were assigned Gene Ontology (GO) Terms based on the SwissProt, PFAM, and/or *D. melanogaster* alignments.

Comparison with 1066 Benchmarking Universal Single-Copy Orthologs (BUSCO v3.0.2, (53)) from arthropods suggests that our transcriptome is of high quality, with 98.4% (91% as single copy and 7.4% as duplicates) of BUSCO genes represented completely and 0.3% represented by fragmented sequences.

### Light induces short-term down-regulation of genes

Consistent with a large and immediate effect of light on gene expression, there were many more differentially expressed genes in the group receiving the light pulse than in the one that did not. This trend was observed at both sampling points, 1 and 10 hours after the light pulse (compare panels C and D in Figure 2; Supplementary File 1: Table S2; Supplementary File 2: Tables S3-S6). Also, consistent with an immediate effect of light, more genes were differentially expressed between the two collection times in the group that received a light pulse than the group that did not (compare panels A and B in Figure 2). The individuals receiving the light pulse sampled 1-hour later show the largest number of differentially expressed transcripts (Figure 2A).

**Figure 2.**
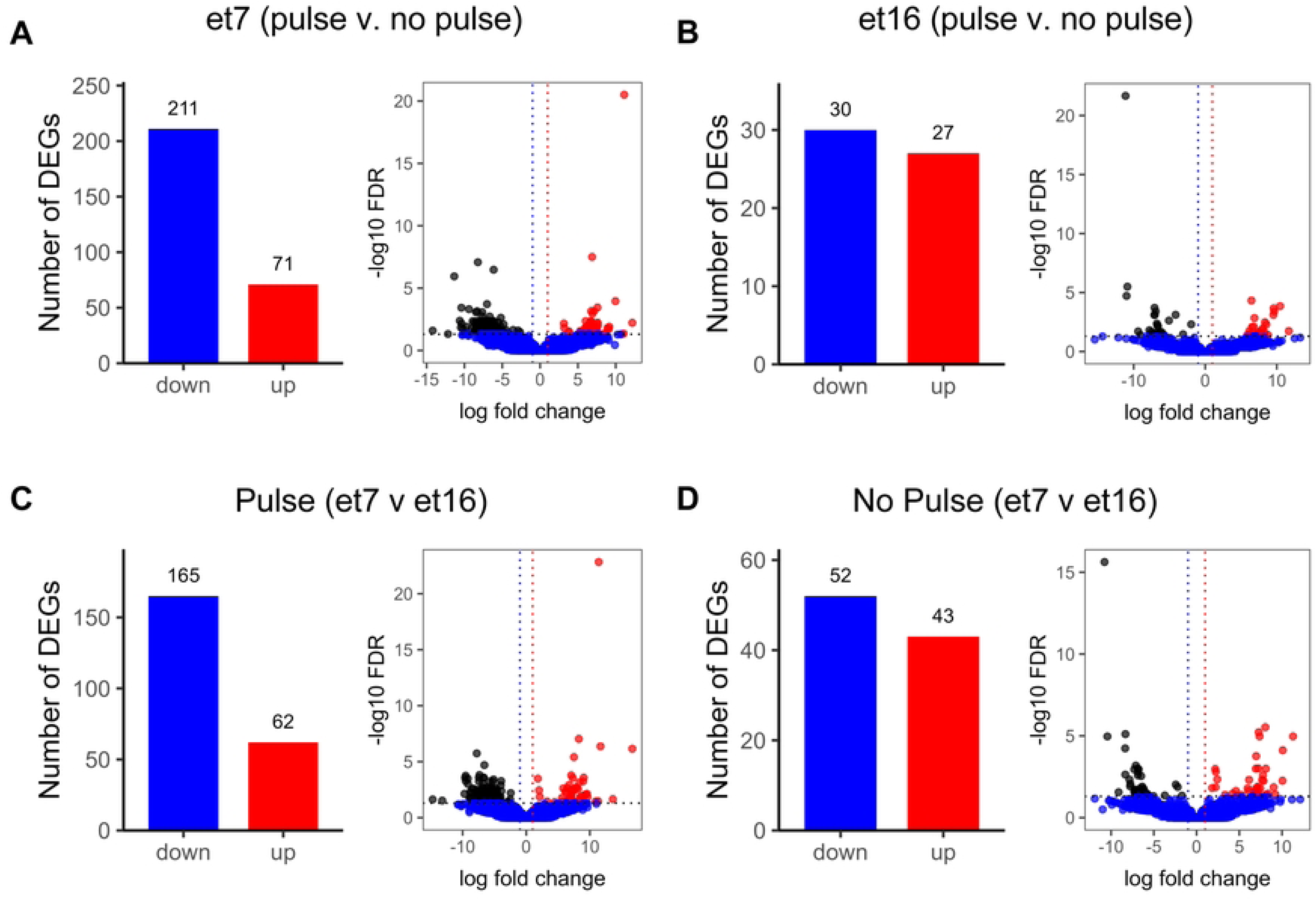
Identification of differentially expressed transcripts associated with light pulse and collection time. Volcano plots showing the negative log10 of the false discovery rate value (Y-axis) against log2 of the fold change (X-axis). The up and down-regulated differentially expressed transcripts at a false discovery rate (FDR) < 0.05 with |log fold change|>1 are depicted in blue and red, respectively. (**A**) The differentially expressed transcripts of spiders given light pulse relative to spiders without light pulse at et7. (**B**) The differentially expressed transcripts of spiders given light pulse relative to spiders without light pulse at et16. (**C**) The differentially expressed transcripts of spiders collected at et7 relative to spiders collected at et16 with a light pulse. (**D**) The differentially expressed transcripts of spiders collected at et7 relative to spiders collected at et16 without a light pulse.

We further found that the light pulse was more likely to down-regulate than up-regulate genes (Figure 2, Supplementary File 1: Supplementary Table S2). The most dramatic shift toward down-regulation was observed at the first collection point, *et7* (Figure 2A), where between light pulse and no light pulse 211 transcripts were down-regulated, but only 71 were up-regulated. Such a large difference in differentially expressed (DE) transcripts might suggest an overall inhibitory effect of a light pulse. However, this inhibitory influence of light disappeared by the time of the second collection, *et16*, when the number of down-regulated genes was very close to that of up-regulated genes (30 vs. 27). Furthermore, for the group experiencing the light pulse, transcript abundance was lower at the early collection time (*et7*) than at the later collection time (*et16*) (Figure 2C, 165 down vs 62 up). In contrast, without the light pulse, similar numbers of transcripts were up- and down-regulated between *et7* and *et16* (Figure 2D, 52 down vs 43 up). These results again suggest an inhibitory effect of light on transcription.

### Flipped patterns of expression identify candidate circadian genes

The 1-hour light pulse applied at *et5* is analogous to the time at which spiders experience the most extreme phase advance and delays of locomotor activity (Fig. 1A); therefore, the light pulse should accordingly shift the phase of expression of circadian genes. We explored how the light pulse might shift the phase of gene expression by modifying two mathematical models, one based on *D. melanogaster* and one based on *Neurospora crassa* (Supplementary File 1). Once we identified parameters that would recapitulate *M. wittfeldae*’s FRP, entrainment profiles, and PRC, we input those parameters to visualize how a 1-hour light pulse would shift the phase. For both models, we found that at *et7,* circadian genes should be downregulated by a light *pulse* compared to the *no pulse* condition (Figure 3A, top). In contrast, at *et16* the pattern of gene expression should be flipped, such that mRNA in the *pulse* condition should be higher than the *no-pulse* condition (Figure 3A). As a result, we expected the ratio of mRNA expression between *pulse* and *no pulse* conditions (log Fold Change (FC), see Methods) to reverse between the two collection times, with negative log FC at *et7* and positive log FC at *et16* (Figure 3A, bottom). To visualize this prediction, we plotted log FC (*pulse/no-pulse*) at *et7* against log FC (*pulse/no-pulse*) at *et16* of DE transcripts. We found a large cluster of DE transcripts in the upper left quadrant, consistent with the flipped pattern of expression predicted by our models (Figure 3; Supplementary File 3: Table S7).

**Figure 3.**
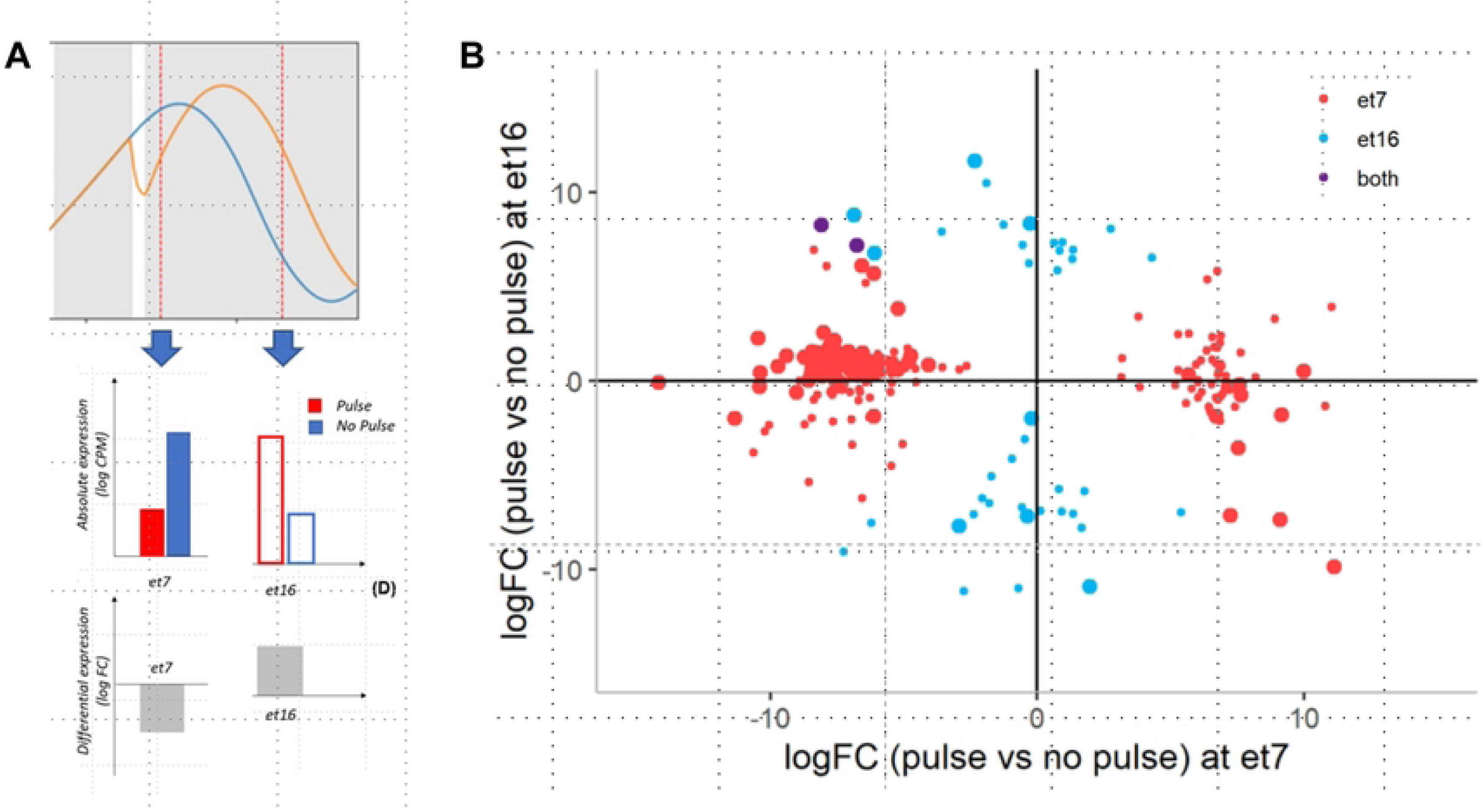
Genes with a shifted phase of gene expression should show a flipped pattern of expression. **(A)** Modeling simulation of the pulse of light. At top – our modified *D. melanogaster* model shows the pattern of mRNA abundance for a clock gene that remains in constant darkness (blue) or receives a 1-hour pulse of light at et5-6 (orange) after entraining to a 12:12 LD cycle (see Supplementary Figure S1). In middle – model-generated absolute gene expression at et7 is higher for no-pulse (red) than pulse (blue) groups; but at et16 the pulse group has higher gene expression than the no-pulse group. Bottom - Differential expression, measured as log of ratio (log FC) between pulse and no pulse conditions, is negative at et7 and positive at et16. **(B)** LogFC of differentially expressed genes (FDR < 0.05, |log FC|>1) between pulse and no pulse condition at et7 versus at et16. There is a large cluster of transcripts with negative logFC at et7 but positive logFC at et16, as predicted for genes with a light-induced phase shift. Transcripts differentially expressed between pulse and no pulse at only et7 are red filled dots while genes differentially expressed at only et16 are colored blue. Two genes are differentially expressed at both et7 and et16, colored purple.

The models additionally predicted that the light pulse should change the pattern of expression between our two time points, such that *et7* expression should be higher than *et16* for the *no-pulse* condition but the reverse should be true for the *pulse* condition. Again, we found a large cluster of transcripts with negative log FC (*et7/et16*) for the *pulse* condition but positive log FC (*et7/et16*) for the *no-pulse* condition (Supplementary File 1: Figure S4, Supplementary File 3: Table S8). These DE transcripts with flipped patterns of gene expression, and thus potentially phase-shifted by light, are enriched for multiple GO terms (Supplementary File 1: Table S9). Some of these enriched GO terms overlapped with ones that were enriched for genes differentially expressed in the 4 pairwise comparisons (Supplementary File 4: Tables S10-S13), especially ones related to cell growth and development.

### Canonical clock genes are present in spiders but are weakly differentially expressed

We identified orthologs of each of 6 canonical clock genes (*tim, per, cyc, clk, cry1, cry2*) in our *M. wittfeldae* transcriptome using a phylogenetic approach (see Methods for details). Only *cry2* was significantly differentially expressed in any of our pairwise comparisons (FDR < 0.01). Specifically, *cry2* was more highly expressed at the second collection time (*et16*) than the first collection time (*et7*) when not subject to a light pulse (Figure 4, Supplementary File 1: Figure S5). The significant difference between time points in the constant dark for *cry2* is consistent with it being a cycling gene, as in butterflies and mammals. The loss of a significant difference between time points when pulsed with light is also consistent with light shifting the phase of *cry2*, as is the flipped pattern of expression between *et7* and *et16* when comparing *pulse* to *no pulse* groups (Figure 4). *Clk* had a similar pattern of expression as *cry2*, although none of the comparisons met the FDR threshold for significance (Figure 4, Supplementary File 1: Figure S5).

**Figure 4.**
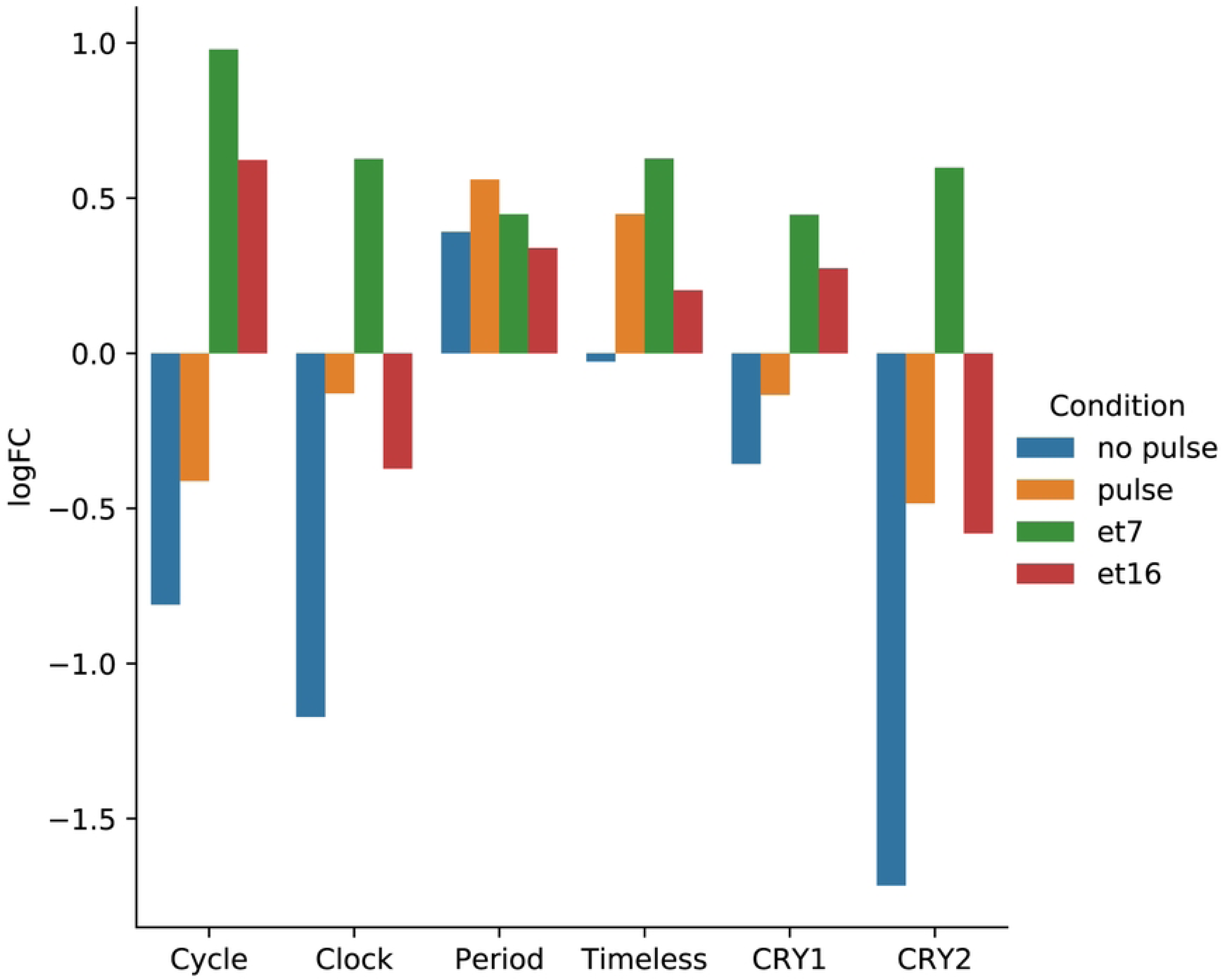
Expression patterns of six canonical clock genes. No Pulse = et7/et16 in the group that remained in darkness; pulse = et7/et16 in the group that received a 1-hour light pulse. et7 = light pulse/no pulse at collection time 1 hour (et7) after the light pulse; et16 = light pulse/no pulse at the collection time 10 hours after the light pulse (et 16).

## Discussion

### First Phase Response Curve for a spider suggests biological mechanisms of their circadian clock

A fundamental property of circadian oscillators is that the internal phase of rhythm is synchronized with the environmental light-dark cycle through the mechanism known as entrainment (54). To describe the underlying dynamics of entrainment, it is useful to perturb a rhythm with a single stimulus (such as a light pulse) and measure the response in subsequent circadian cycles. This approach, pioneered by Pittendrigh, is described by a nonparametric (discrete) model of entrainment (27). The model assumes that light at discrete times of day (e.g., dawn and dusk) falls at specific phases of the pacemaker and causes instantaneous phase shifts that are equal to the difference between the period of the rhythm and the period of the LD cycle. The phase response curve (PRC), which is a graphical representation of the shifts in response to stimuli given at different phases, have been extensively used to quantify nonparametric entrainment mechanisms (55,56). Numerous experiments have described PRCs in multiple model organisms (56) with remarkable similarities in the pattern observed: the response to light typically is largest at time points where the organism is not exposed to the light under entrained conditions. In this regard, our experimental PRC demonstrates a similarity to other models described so far: the largest phase shift in response to light occurred during the subjective night, and a “dead zone” was observed during the subjective day.

Two types of PRC can be distinguished: weak resetting PRC (type 1) and strong resetting PRC (type 0). The type 1 PRC has small (typically less than two hours) phase shifts, while type 0 has larger phase shifts (57). The PRC is considered a property of the circadian pacemaker, and its type depends on the strength of, or sensitivity to, the stimulus (58–60). It has been demonstrated in several model organisms that an increase in light intensity or light duration can induce a PRC transition from type 1 to type 0 (61). It is, therefore, surprising that our experiment reveals a strong, type 0, PRC in response to a brief (1 hr.) stimulus of average intensity (1,400 lux light is comparable to sunlight in the early afternoon of an overcast day). In insects, such light intensity typically results in type 1 (weak) PRC. For example, in cockroach *Nauphoeta cinerea*, ∼1,600 lux stimulus of 3 hours duration leads to a weak type 1 PRC, and an increase in stimulus duration to 12 hours is needed to switch to type 0 (62). In the mosquito *Culex pipiens quinquefasciatus*, 2-hour 8,000 lux pulses were needed for transition to type 0 (63). For *D. melanogaster*, even 12 hours of 3000 lux light pulse still induces type 1 PRC, with the transition to type 0 only possible in *per* mutants (37).

The strength of the light stimulus is not the only potential mechanism to induce a strong, type 0, PRC. The theory of oscillation states that the balance between the stimulus intensity and oscillator amplitude is crucial for PRC shape. That is, the low amplitude (weak) oscillator can produce a large change in response to a moderate stimulus (39). Therefore, it is possible that if spiders’ clock proteins have low amplitude oscillation, a light pulse of normal amplitude can produce large changes. A low amplitude oscillator could also explain the limited difference in expression levels between time points for the canonical clock genes (Figure 4, Supplementary File 1: Figure S5).

Another possible explanation for the observed type 0 PRC in *M. wittfeldae* might be that light is a very strong stimulus for spiders. In other words, the effect of a single photon of light on a circadian protein in spiders could be more powerful than in insects. In this case, spiders would have evolved a unique pathway by which light affects the circadian clock. For example, if the circadian clock is located directly in the eyes, light can affect the clock proteins directly.

Examples of entrainment through the eyes have been documented in birds and mollusks (64,65). In the case of the Japanese quail, the phase response curve is also type 0 (66), suggesting that an ocular circadian clock could increase sensitivity to light. Light could also affect circadian proteins expressed in the brain directly if the carapace is transparent. For instance, *cry1* is expressed in the *D. melanogaster* brain and is directly modified by light (67). We do not currently have evidence that the *M. wittfeldae* carapace is transparent, but circadian expression in the eyes and/or increased sensitivity of a circadian protein like CRY1 to light could contribute to the observed strength of phase shifting in this species.

Since the activity of *M. wittfeldae* (47) and other spider species (46,48,49) increases dramatically at the onset of darkness, light likely has an inhibitory effect on locomotor activity. Inhibition could be direct (e.g. masking) or act on a step in the circadian molecular clock.

Regardless of the neural mechanisms of circadian light response in *M. wittfeldae*, the type 0 PRC is informative about spider entrainment and the potential reasons for the wide variation in FRP among spiders. Since most organisms, including spiders, live in a cycling environment, the phase of entrainment rather than the free-running period should be the main subject of natural selection (68). The endogenous clock may have evolved to time biological events relative to the external day-night cycle. Thus, the phase angle of entrainment, the fixed time interval between the zeitgeber and the internal clock, could be the trait subject to natural selection (69). Our previous findings demonstrate that *M. wittfeldae* and two other spider species have sharp activity onset within 30 minutes of lights off while having a wide range of FRP (46–48). The sharp PRC observed in *M. wittfeldae* and consistent phase angle of entrainment suggest that spiders easily entrain to light, and thus, there may be weak evolutionary pressure to keep FRP close to 24 hours.

### Spider circadian clock gene homologs have unusual patterns of expression

We identified homologs of five genes (*per*, *tim*, *clk*, *cyc*, *cry2*) involved in the transcription-translation feedback loop that appears to be conserved across animals (14–16). We also identified a homolog of the photoreceptor encoding gene (*cry1*) that is primarily responsible for the entrainment of the fruit fly clock. In *M. wittfeldae*, we found *cry2* was significantly differentially expressed between our two sampling times when spiders were not subject to a pulse of light. Interestingly, the pattern of expression was flipped when pulsed with light – e.g. *cry2* was higher at et7 than et16 in constant dark, but lower when pulsed with light (Figure 4). Thus, spiders may retain the ancestral animal clock mechanism in which CRY2 interacts with PER (70,71).

However, patterns of expression for *cry2* and other clock gene homologs suggest spiders may have diverged considerably from the ancestral animal clock. First, none of the other clock gene homologs were differentially expressed between time points. In butterflies and mammals, CRY2 interacts with PER to repress their own transcription, and the expression patterns of *cry2* and *per* are synchronized (70,72–74). In *M. wittfeldae*, however, *per* expression was consistent across all conditions rather than mirroring the expression pattern of *cry2* (Supplementary Figure S3). The clock gene homolog with the most similar expression pattern to *cry2* was instead *clk*, which shows the same pattern of lower expression at et7 than et16 (*P* = 0.0069, Supplementary File 1: Figure S3), although its FDR did not meet the threshold for significance. In all examined insects and in mammals, CLOCK promotes transcription of *per* and other clock genes when it dimerizes with CYCLE/BMAL1. In fruit flies, *clk* is rhythmically expressed, but with two peaks per day (10), and in honeybees, *clk* expression mirrors the cycling of *per*, *tim*, and *cry2* (70). On the other hand, many insects and mammals have a constant expression of *clk* throughout the day, and instead, *cyc* is rhythmically expressed (70,75,76). We found *cyc* was very lowly expressed (< 1 TPM) in all our conditions. Further experiments are needed to determine if this low-level expression produces sufficient CYCLE to dimerize with CLOCK or if CLOCK can promote transcription on its own or with a different partner in spiders.

In fruit flies and potentially other insects, PER interacts with TIM to repress their own transcription and have synchronized expression (73,77,78). We found that *tim*, similar to *per*, has a pattern of near-constant expression levels across time and light pulse conditions. It is possible that these two proteins interact in spiders and are involved in the clock’s transcription-translation feedback loop, but that our two time points happened to coincide with rising and falling phases rather than the peak and trough. It is also possible that the amplitude of oscillations is low, which would be consistent with the type 0 PRC for *M. wittfeldae*, since low amplitude oscillators are easily perturbed (40,79,80). However, sampling more times points is needed to establish the rhythmic, or lack of rhythmic, expression of these conserved clock genes.

We were especially interested in discovering the photoreceptor encoding homolog, *cry1*, since spiders easily entrain to light:dark cycles and CRY1 is essential for robust entrainment in fruit flies (25,67,81) and other arthropods (72,82). *Cry1* transcript abundance did not differ between time points or light pulse conditions. In fruit flies, *cry1* cycles, but light does not directly impact transcription levels of *cry1*. Instead, light changes the conformation of CRY1 to cause degradation of TIM, so it may not be surprising that our pulse of light did not affect the transcript abundance of *cry1*. Like *per* and *tim*, our two sampling times may not have caught the peak and trough of *cry1* expression, *cry1* may not oscillate, or the amplitude of oscillations may be low. Further experiments are needed to determine if *cry1* encodes a photoreceptor in spiders and if it is involved in circadian entrainment.

### Light downregulates many genes and may shift the phase of gene expression

When we applied a light pulse at the time of night when *M. wittfeldae* showed the greatest behavioral sensitivity to light (Figure 1), we found many more genes were downregulated than upregulated 1-hour after the end of the light pulse (Figure 2). We also predicted that this light pulse would shift the phase of gene expression, resulting in flipped patterns when the light pulse was applied versus not applied (Figure 3). These are potentially cycling genes with a shifted phase due to the light pulse. Admittedly, since we only sampled at 2 time points, there could be other explanations for the flipped patterns of gene expression than a shifted phase. These transcripts had multiple functions and were distributed throughout the cell (Supplementary File 1: Table S9). GO-terms involved in immune response, growth, development, and reproduction were enriched in this candidate set of light-sensitive circadian regulated genes. It has been shown in *D. melanogaster* that the immune response is under circadian control (83,84). Exposure to night light also leads to reductions in immune function in vertebrates and invertebrates (85,86).

We further found enriched GO-terms for growth, development, and reproduction in our candidate phase-shifted genes (Supplementary File 1: Table S9). These results are consistent with prior evidence that exposure to light at night alters circadian rhythms, inhibits reproduction, accelerates development, and leads to increased mortality in vertebrates (85), fireflies (87), and *D. melanogaster* (88). In an Australian nocturnal araneid spider, exposure to artificial light at night accelerated juvenile development, resulting in spiders progressing through fewer molts (89). Light intensity also controls the eclosion circadian rhythm in *D. melanogaster* (90). Our results for *M. wittfeldae* further support a role of light in affecting immune function, development, and reproduction.

## Conclusions

Our results represent the first analysis of a non-insect arthropod Phase Response Curve, which has a strong type 0 shape. The intense phase-shifting response to light could be due to a weak circadian oscillator or to the heightened sensitivity of circadian proteins to light. Patterns of expression of clock gene homologs in spiders may be distinct from other arthropods, and further experiments on these and additional potential cycling proteins could help distinguish the mechanism of spiders’ wide range of FRP and strong entrainment to light. Our transcriptome-wide analysis of expression patterns may represent light-shifted phases of expression of numerous genes, possibly including those directly responsive to light as well as those directly or indirectly influenced by the circadian clock. Thus, we have revealed candidate genes for future research into the spider circadian system.

## Methods

### Experimental Phase Response Curve

Adult female *M. wittfeldae* were wild caught in Washington and Sullivan Counties in northeastern Tennessee, USA in June, July, and August 2017. Individual spiders were placed in clear plastic cups in the laboratory, fed crickets, and lightly sprayed with a mist of distilled water. The spiders were exposed in the laboratory for 2-3 days to an LD 12:12 hr cycle (i.e., 12 hr of light alternating with 12 hr of dark) with the light-dark transition set at 20:00 hr local time. Individual spiders were then placed in clear glass tubes (25 mm diameter x 15 cm length), and the tubes were inserted into locomotor activity monitors (model LAM 25, Trikinetics Inc., Waltham, MA, USA). The monitors were then placed within a temperature-controlled (24 ± 0.5 °C) environmental chamber. The activity was recorded continuously for each spider in 1-minute bins via interruptions of infrared beams in the activity monitors.

Using an Aschoff type 1 protocol (91), a phase response curve was derived from the results of four trials conducted from June to September 2017. Spiders in the monitors were exposed to

5-8 continuous days of constant dark (DD) prior to a 1-hour light pulse (approximately 1400 lux) and another 6-11 full days of DD after the light pulse. The pulses, provided by four vertically mounted fluorescent tubes, were delivered at a different time of day in each trial. The onset of the pulse was considered the stimulus phase. Because of the broad range of free-running periods (FRPs) in this species (Jones et al. 2018), the pulses delivered in the four trials cumulatively encompassed the entire span of the circadian cycle. In two trials, the spiders transitioned directly from the laboratory LD cycle to DD conditions. In the other two trials, the spiders were given an additional two or three complete days of LD 12:12 hr (with the light-to-dark transition at 20:00 hr, as in the other trials) in the monitors before transitioning to DD.

The activity data were displayed as double-plotted actograms for each spider using ClockLab Analysis 6 Software (Actimetrics, Wilmette, IL, USA). To determine the circadian time (CT) at which the light pulse occurred for individual spiders, activity onsets were used as the phase reference points, with activity onset denoted as circadian time, CT 12. This is consistent with nocturnal organisms that become active at “dusk.” *M. wittfeldae* is highly nocturnal with locomotor activity beginning shortly after the light-to-dark transition (47). Phase shifts were calculated by tracking the pre- and post-pulse FRPs using the linear regression function in ClockLab through activity onsets for each individual spider. A t-test for correlated samples was used to determine if there was a significant change in FRP following the light pulse.

### Gene Expression Experimental Design

Based on the shape of the PRC, we applied a 1-hour light pulse at the point in time when *M. wittfeldae* showed maximal phase advances and delays. We used two mathematical models (detailed in Supplementary File 1) to predict collection time points that would maximize the difference in gene expression levels between spiders receiving the light pulse and those that did not. We also selected two collection time points after the light pulse to maximize the ratio of mRNA between time points for each experimental group.

Adult female *M. wittfeldae* were collected on 16 July 2018 in Washington County, TN, and placed in clear plastic deli jars on a lab bench receiving natural lighting. All spiders were provided with a wet cotton ball and offered a small cricket. On 18 July 2018, each spider was moved to a clear glass or plastic tube (25 mm diameter x 15 cm length) and inserted into a locomotor activity monitor (model LAM25, Trikinetics Inc., Waltham, Massachusetts, U.S.A.). The monitors were placed into two separate temperature-controlled (24 ± 0.5 °C) environmental chambers with the lights on (1400-1600 lux) at ∼16:00 until 20:00, after which the spiders remained in the dark for 3.5 days (84 hours) to establish the free-running period of each individual. In order to synchronize the circadian rhythms of the individual spiders, they were then entrained to a 12:12 LD cycle for five days. On the evening of the fifth day, the lights went out at 20:00 (*et0*), as per the prior 4 days. However, one chamber received 1 hour of light (1400-1600 lux) starting 5 hours after the lights were turned off (*et5-et6*, 01:00-02:00 27 July 2018) (*pulse* group). The other chamber remained dark (*no pulse* group). One hour after the end of the light pulse (*et7*, 03:00) 5 spiders were removed from each environmental chamber and snap frozen in liquid nitrogen under dim red light. Both chambers remained dark until the final collection time 9 hours later (*et16*, 12:00), when five spiders were again removed from each chamber and snap frozen in liquid nitrogen under dim red light.

### RNA-sequencing, transcriptome assembly, and annotation

Total RNA was isolated from one half of a cephalothorax (the fused head-body) of each of the 20 experimental spiders by homogenizing tissue in TRIzol (Invitrogen) and further purifying with the RNeasy Mini Kit (Qiagen) with on-column DNaseI digestion to remove contaminating DNA. RNA quality and quantity were verified with the Agilent 2100 Bioanalyzer, and 20 individually barcoded cDNA libraries were constructed with the TruSeq kit (Illumina) by the Genomics Research Laboratory at the Biocomplexity Institute, Virginia Tech. All 20 libraries were multiplexed and sequenced in a single lane with the NextSeq 500 (Illumina) using the High Output mode with 300 cycles (150 base paired end reads). De-multiplexing and barcode removal were performed at Virginia Tech.

Transcripts were *de novo* assembled with Trinity v2.8.4 (92) from all 20 RNAseq libraries. Raw reads were trimmed of low-quality base calls using Trimmomatic (93), and *in silico* normalization was used to reduce the number of reads entering the assembly phase, using scripts in Trinity.

Annotation involved a multi-step process using the Transcriptome Trimming and Annotation Pipeline (TrTAP) (94). Trimming started when any transcripts with a significant BLASTN match to the SILVA ribosomal database v.132 (95) or a tRNAScan v.1.3.1 match (96) were identified (RNA) and removed from further analysis. Second, all the transcripts were compared to a custom-made database of spider silk proteins encoded by the spidroin gene family (97) and subsequently to the proteomes of 5 arthropod species using BLASTx. These species included 3 spiders (listed from most closely related to least): *Araneus ventricosus* (GCA_013235015.1_Ave_3.0, Kono et al. 2019)), *Trichonephila clavipes* (MWRG00000000.1, (99)), and *Stegodyphus mimosarum* (GCA_000611955, (100)); another arachnid, *Ixodes scapularis* (GCF_016920785.1,Gulia-Nuss et al. 2016); and an insect, *D. melanogaster* (FB2019_08 (102)). The *D. melanogaster* gene matches with a cutoff of 1e-40 was used to identify chimeric sequences with a custom Python script described in (103).

For each Trinity “gene”, the transcript with the best BLAST alignment, as determined by the highest bit-score, or with the longest open reading frame (ORF) if there were no BLAST alignments, was chosen to represent that gene, with all the genes used to calculate the expression with RSEM (104). To further reduce redundancy and identify fragments of genes, the best matching gene of each protein in the curated silk gene database plus the 5 species proteomes were also identified. These reciprocal best matches were always retained for downstream analyses (BEST, Supplementary File 1: Table S1). If a Trinity gene was not the best match of a database protein, it was considered “GOOD” if it aligned to >20% of the database protein and exceeded 1 TPM (transcript per million) in at least one RNAseq library. Genes that aligned to >20% of the database protein and did not meet the 1 TPM threshold were dubbed “LOW_EXP” (Supplementary Table S1). Genes that aligned to <20% of the database proteins were dubbed “LOW_COV” (Supplementary Table S1). For the genes with no BLAST alignments to a proteome, the gene was called “LONGORF” if the ORF exceeded 50 amino acids and had >1 TPM in at least one RNAseq library (Supplementary Table S1). ORFs that exceeded 50 amino acids but had <1 TPM were called “LOW_EXP_LONGORF” (Supplementary Table S1). Genes that did not have a BLAST hit and an ORF of less than 50 aa were called “NO_HIT”. A reduced set of probable protein-coding genes (BEST, GOOD, LONGORF, and LOWEXP) was then translated using the direction and frame of the matching BLASTX hit or, in cases without such hit, translated according to the longest open reading frame. These translated genes were further annotated by comparison to SwissProt with BLASTP and to PFAM with HMMER v 3.2.1 (105) with Gene Ontology (GO) terms (106,107) assigned to each “gene” based on the best alignment to *D. melanogaster*, SwissProt, and PFAM, with GO SLIM annotations obtained from GO SLIM viewer (108). The completeness of the reduced transcriptome was assessed with the set of single-copy orthologous genes in arthropods (BUSCO v3, (53)).

### Canonical circadian gene identification

We used a phylogenetic approach to determine which transcript was an ortholog of the canonical clock gene in *D. melanogaster* or *Apis mellifera*: *clk*, *cyc/bmal1*, *tim*, *per*, *cry1*, *cry2*. We first used Orthofinder v2.4.1 (109) to identify groups of homologous genes among 6 spider species, a tick, and two insects. In addition to our *M. wittfeldae* transcriptome, we used the predicted proteomes from published transcriptomes or genomes for the following species: *Araneus ventricosus* (Araneidae, BGPR01000000.1), *Trichonephila clavipes* (Araneidae, MWRG01000000.1), *Latrodectus hesperus* (Theridiidae, GBJN01000000.1), *P. tepidariorum* (Theridiidae, GCF_000365465.3), *Stegodyphus mimosarum* (Eresidae, GCA_000611955.2), *I. scapularis* (Ixodidae, GCF_016920785.1), *D. melanogaster* (Drosophilidae, FB2019_08), *A. mellifera* (Apidae, GCF_003254395.1). We then identified the “orthogroups” containing the canonical circadian gene of *D. melanogaster* or *A. mellifera* (Supplementary File 1: Table S2). For 5 of the genes, only a single *M. wittfeldae* transcript was placed in the orthogroup; these were considered the ortholog of the circadian gene. For the cryptochromes (*cry1* and *cry2*), we inferred a phylogenetic tree using maximum likelihood (RAxML,(110)). From the resulting tree, we chose the single *M. wittfeldae* transcript in the clade with the canonical circadian clock gene (Supplementary File 1: Figure S6).

### Differential gene expression analysis

Differential gene expression among experimental groups was based on gene-level expression estimates derived from transcript abundance estimates as recommended by Soneson et al. (111). In brief, the expected read counts of each Trinity-assembled transcript were calculated by RSEM v.1.3.1 (104), which takes into account read-mapping ambiguity due to multiple isoforms or even alleles having been assembled. The gene-level counts were then calculated as the sum of the included transcripts, weighted by their length as described in Soneson et al. (111) and implemented through Trinity v2.8.4. Because RSEM can distribute one read among multiple transcripts, some counts were not integer values. These values were rounded for input into differential expression analyses, which require integer values for read counts. Prior to further expression analyses, genes unlikely to encode proteins were removed (Supplementary Table S1). For comparing expression levels among circadian genes, we used Transcripts Per Million mapped transcripts (TPM) normalized for differences in sequencing depth among libraries with the Trimmed Mean of M (TMM) values calculated with EdgeR (112).

Four different pairwise gene expression analyses were performed on the read count data using the EdgeR Bioconductor package (112): 1) comparing samples with light pulse to samples without light pulse at et7; 2) comparing samples with light pulse to samples without light pulse at et16; 3) comparing samples collected at et7 to samples collected at et16 with a light pulse; 4) comparing samples collected at et7 to samples collected at et16 without a light pulse. A gene was kept in the analysis if it had sufficient expression (TPM > 0.1) in at least two samples. The raw library sizes were normalized using the TMM method from EdgeR. The significantly differentially expressed genes in the pairwise comparisons were identified with both a false discovery rate (FDR) < 0.05 and an absolute value of log FC > 1 (Supplementary File 2: Tables S3-S6).

### Gene Ontology analysis

GO analysis was conducted to determine the functions enriched in DE genes relative to all annotated genes using the ‘GOseq’ Bioconductor package (113) based on Wallenius non-central hyper-geometric distribution, with a p-value of 0.1. The lists of GO terms, along with their p-values generated from ‘Goseq’, were summarized and visualized by the REViGO online tool (114) (Supplementary Figure S7).

## Acknowledgements

We thank Jess Petko for conversations about clock genes. This work was funded by the National Science Foundation (IOS-2235711 to NT and NAA, IOS-2235710 to DM and TCJ).

## Supplementary Materials

**Supplementary File 1**. Description of two mathematical models to predict shifts in gene expression in response to light. The file also includes Supplementary Figures S1-S7 and Supplementary Tables S1, S2, S9, and S14.

**Supplementary File 2**. Includes Supplementary Tables S3-S6, which describe the differentially expressed transcripts identified in each of the four pairwise comparisons.

**Supplementary File 3**. Includes Supplementary Tables S7-S8, which describe the differentially expressed transcripts with flipped log FC.

**Supplementary File 4**. Includes Supplementary Tables S10-S13, which describe the GO terms enriched for differentially expressed transcripts.

## Data Availability

Raw sequence reads and assembled transcriptome are available in NCBI through Bioproject PRJNA767861. Additional data are reported in the manuscript, the supplementary files, and https://github.com/Toporikova-Lab/Spider-RNA-seq-analysis/tree/c2707f13824e38e1498329e3f7591111863385a1/Modeling

